# Microglia play beneficial roles in multiple experimental seizure models

**DOI:** 10.1101/2023.03.04.531090

**Authors:** Synphane Shelton-Gibbs, Jordan Benderoth, Ronald P. Gaykema, Justyna Straub, Kenneth A. Okojie, Joseph O. Uweru, Dennis H. Lentferink, Binita Rajbanshi, Maureen N. Cowan, Brij Patel, Anthony Brayan Campos-Salazar, Edward Perez-Reyes, Ukpong B. Eyo

**Author notes:** Correspondence: Ukpong B. Eyo, Ph. D, Assistant Professor, Center for Brain Immunology and Glia, Neuroscience Department, University of Virginia School of Medicine.

## Abstract

Seizure disorders are common, affecting both the young and the old. Currently available antiseizure drugs are ineffective in a third of patients and have been developed with a focus on known neurocentric mechanisms, raising the need for investigations into alternative and complementary mechanisms that contribute to seizure generation or its containment. Neuroinflammation, broadly defined as the activation of immune cells and molecules in the central nervous system (CNS), has been proposed to facilitate seizure generation, although the specific cells involved in these processes remain inadequately understood. The role of microglia, the primary inflammation-competent cells of the brain, is debated since previous studies were conducted using approaches that were less specific to microglia or had inherent confounds. Using a selective approach to target microglia without such side effects, we show a broadly beneficial role for microglia in limiting chemoconvulsive, electrical, and hyperthermic seizures and argue for a further understanding of microglial contributions to contain seizures.

## 1. INTRODUCTION

Seizure disorders affect 65-70 million individuals globally. Prolonged seizures, known as status epilepticus, can cause significant brain alterations and even damage. Notably, seizures are often self-limiting and understanding endogenous mechanisms by which seizures develop and lead to chronic epilepsy can offer insights into developing promising therapeutic approaches. Development of antiepileptic drugs (AEDs) have been based on targeting neuronal mechanisms of excitability. However, while neurons are central to the development of seizures, neurons do not function independent of other cells and other cellular factors. Indeed, a network of glia, traditionally considered the support cells of the CNS, are critical for proper neuronal development and function (Allen & Lyons, 2018; Thion, Ginhoux, & Garel, 2018). Interestingly, there is growing evidence that glia contribute to epileptic phenotypes (Eyo, Murugan, & Wu, 2017; Robel & Sontheimer, 2016).

Inflammation has been implicated in both sterile and infection-associated seizures (Devinsky, Vezzani, Najjar, De Lanerolle, & Rogawski, 2013; Vezzani, 2014; Vezzani, Aronica, Mazarati, & Pittman, 2013) including paraneoplastic encephalitis (Serafini et al., 2016) and Rasmussen encephalitis Rasmussen encephalitis (Cay-Martinez, Hickman, McKhann Ii, Provenzano, & Sands, 2020). For example, mice lacking interleukin-1 receptor (IL1R1), the receptor to IL-1α and IL-1β, are more resistant to seizures (C. Dube, Vezzani, Behrens, Bartfai, & Baram, 2005). Notably, IL-1β infusions enhanced seizures, while infusions of IL-1ra, an endogenous IL-1 antagonist mitigated seizures (Heida & Pittman, 2005). Similarly, TNF-α infusions in neonatal rats facilitated chemoconvulsive seizure development in adulthood, while treatment of mice with function blocking TNF-α antibodies reduced experimental seizures in adulthood (Galic et al., 2008). Moreover, interferon-alpha (IFN-α) has been implicated in febrile seizures (Masuyama et al., 2002). Thus, inflammatory cytokines like IL-1α, IL-1β, TNF-α and IL-1α facilitate the initiation of seizures and subsequent progression to epilepsy.

Despite the above observations, inhibiting inflammation during seizures has yielded conflicting results. For example, COX inhibitors failed to reduce epileptogenesis (Holtman, van Vliet, Edelbroek, Aronica, & Gorter, 2010; Holtman et al., 2009) and dexamethasone (DEX), a broad acting anti-inflammatory agent, gave conflicting results in seizure severity (Al-Shorbagy, El Sayeh, & Abdallah, 2012; Duffy, Chun, Ma, Lythgoe, & Scott, 2014; Marchi et al., 2011). Moreover, cytokines are sometimes significantly elevated (Haspolat et al., 2002; Virta, Hurme, & Helminen, 2002), unchanged (Lahat, Livne, Barr, & Katz, 1997), or partially elevated (Ichiyama, Nishikawa, Yoshitomi, Hayashi, & Furukawa, 1998) in children following febrile seizures, raising questions as to the power of the cytokine hypothesis in explaining seizure generation. Poignantly, inflammation can be induced by different cell types including astrocytes, microglia, and peripheral cells that may differ in their actions. For example, IL-1β is principally produced by astrocytes and not microglia during seizures (C. M. Dube et al., 2010). Therefore, broadly targeting inflammation may mask distinct roles employed by each of these cells during seizures raising a need for more selective cell-type specific targeting.

Microglia are the primary brain-resident inflammation-competent cells. Conflicting experimental results suggest that microglia either promote seizures (Abraham, Fox, Condello, Bartolini, & Koh, 2012; Di Nunzio et al., 2021; Eun, Abraham, Mlsna, Kim, & Koh, 2015; Kim et al., 2015) or dampen neuronal hyperactivity (Cserep et al., 2020; Kato et al., 2016; Y. Li, Du, Liu, Wen, & Du, 2012; Merlini et al., 2021) and perform seizure-limiting functions as evidenced when they are eliminated by pharmacogenetic approaches (Mirrione et al., 2010; Wu et al., 2020). However, concerns about these pharmacogenetic elimination approaches have arisen with evidence that they cause disrupted structural brain features, behavioral, and cellular abnormalities, as well as increased cytokine release and glial reactivity (Bedolla et al., 2022; Rubino et al., 2018). These results confound the interpretation of previous findings as to the roles microglia play in seizures. Moreover, the extent to which microglia regulate seizure phenotypes have been explored mainly in chemoconvulsive seizure paradigms while their roles in alternative seizure paradigms have remained largely unexplored. In the current study, we used a pharmacological approach that eliminates brain microglia without altering either cytokine levels, glial reactivity, or affecting behavior. We then applied this approach in multiple seizure models and provide compelling results to argue for beneficial roles for microglia in limiting seizures.

## 2. MATERIALS AND METHODS

### 2.1 Animals

All animal experiments were carried out pursuant to the relevant guidelines and regulations of the University of Virginia and approved by the Institutional Animal Care and Use Committee with protocol number 4237-08-21. The animals were housed under controlled temperature, humidity, and light (12:12 hr. light: dark cycle), with food and water readily available ad libitum. This study used both male and female mice on a C57BL/6J background between 6-12 weeks of age, and consisted of the following genotypes: CX3CR1^GFP/+^ expressing GFP under control of the fractalkine receptor (CX3CR1) promoter (Jung et al., 2000); and CX3CR1^Cre/+^:Rosa26^iDTR/+^ mice (Buch et al., 2005; Parkhurst et al., 2013); VGAT-Cre mice (Vong et al., 2011); TRAP2 mice capable of expressing tamoxifen-inducible tdTomato under the control of cFos promoter (DeNardo et al., 2019) and C57Bl/6J mice as wildtype mice. Mice were housed in groups for the experiment without special environmental enrichment.

### 2.2 Treatments for microglial elimination

#### 2.2.1 Pharmacogenetic elimination

Both CX3CR1^Cre/+^ and CX3CR1^Cre/+^:Rosa26^iDTR/+^ mice received two doses of diphtheria toxin (DT, 50 μg/kg) by intraperitoneal injection at 48h intervals and seizure experiments were conducted 24 h after the last DT injection.

#### 2.2.2 Pharmacological elimination

Microglia were depleted using a potent CSF1R inhibitor, PLX3397, that has been previously shown to lack the inflammatory consequences during the elimination process. Microglia from the adult brain were depleted by feeding adult mice with chow containing PLX3397 (660 mg/kg) for 7 days. Similarly, for febrile seizure studies, pre-weaned P10-P12 pups were administered 50μl of PLX3397 (40 mg/kg) or vehicle (10% DMSO; 50% polyethylene glycol; 40% saline) by daily intraperitoneal (i.p.) injections from P10-12. For studies using TRAP2 mice to label hyperactive neurons at the desired time points, tdTomato expression coupled to cFos expression was induced by administering mice with a single intraperitoneal (i.p.) injection of tamoxifen (40 mg/kg) dissolved in corn oil. Mice were allowed to return to their home cages for another week, after which they were perfused, and used for further imaging and analysis.

### 2.3 Seizure induction

#### 2.3.1 Chemoconvulsive seizure induction

For chemoconvulsive seizure initiation model, 2-4-month-old mice were used, and i.p. injected with kainic acid (KA) at 24-27 mg/kg body weight. Control mice were injected with equal volume of saline that was used as the vehicle to dissolve the kainic acid. Seizures were scored using a modified Racine scale as follows: (1) freezing behavior; (2) rigid posture with raised tail; (3) continuous head bobbing and forepaws shaking; (4) rearing, falling, and jumping; (5) continuous occurrence of level 4; and (6) loss of posture and generalized convulsion activity (Avignone, Ulmann, Levavasseur, Rassendren, & Audinat, 2008; Eyo et al., 2014; Racine, 1972). Scores were recorded every 5 minutes for up to 4 hours. Scores were given based on the summary of the mouse behavior over the 5 minute interval. The experimenter was “blind” to the prior condition of the mice. The seizure scores are presented as the median of the scores and the area under the curve (AUC) was analyzed.

#### 2.3.2 Febrile seizure induction

For hyperthermia-induced febrile seizure induction, P13-P15 mice pups were placed in a fiberglass hyperthermia chamber heated by means of an overhead heat lamp and a basal heating plate. Temperatures of the two heat sources was precisely regulated by a temperature regulator. In experiments measuring the threshold temperature to seizure initiation, the body temperature of the pups was monitored using a rectal temperature probe connected to a digital thermometer. Baseline activity of mice was first recorded for 15-30 minutes at normothermic conditions 35°C, following which hyperthermia was induced by subjecting them to temperatures around 41-43°C, for 30 minutes, or until seizures were induced. Video monitoring and recording of the seizure induction was carried out. Latency to first signs of inactivity and convulsions, and temperature threshold for seizure initiation were all recorded and used to compare seizure outcomes between experimental and control groups.

### 2.4 EEG implantation, recordings, and electrical stimulation protocols

Surgeries were performed on 8-12 week-old mice using isoflurane anesthesia. Ketoprofen was used as an anesthetic for the surgery. An EEG recording headset included bipolar depth electrodes for electrical stimulation composed of Teflon-coated stainless-steel wire (A-M Systems, diameter = 0.008”, #791400). Electrodes were soldered to a plastic pedestal (Plastics One) and secured to the skull with dental cement. A Kopf stereotaxic apparatus was used to place the depth electrodes (from bregma): 3 mm posterior, 3 mm lateral, and 3 mm depth.

For EEG recording, mice were connected to a video-EEG monitoring system (AURA LTM64 using TWin software, Grass) via a flexible cable and commutator (Plastics One). One day later, the after-discharge threshold (ADT) was measured. The hippocampal depth electrode was connected to a constant current stimulator (A-M Systems, Model 2100), which delivered a 2 s train of 1-ms pulses at 50 Hz. The current was initially set at 10 μA and increased in 10-μA increments until an electrographic discharge was observed (20-120 μA).

For kindling of VGAT-Cre mice, the current intensity was set to 1.5 × the magnitude of ADT for that mouse and was delivered six times per day separated by at least 1 hour. Animals were considered fully kindled when stimulations evoked five consecutive seizures that triggered 5 motor seizures with bilateral clonus and loss of posture control. Animals were monitored by video/EEG throughout the experiment (24 h/d, 7 d/wk). Spontaneous seizures were defined as high amplitude spike-wave discharges with >2-Hz frequency lasting at least 15 s. Electrographic seizures were verified by examining the corresponding video and a behavioral score was assigned. All seizures reported in this study had a motor component.

Similar methods were used for continuous hippocampal stimulation (CHS) including electroencephalography and behavior patterns during experimental status epilepticus as detailed previously (Lewczuk et al., 2018). These studies used C57BL/6J mice obtained from Jackson Labs (N=32). Only males were used since female reproductive cycles affect seizure frequency (Joshi et al., 2018). The electrical stimulation protocol is similar to kindling, but the duration is increased to 10s and the total train duration is 15 s. The 5 s off interval is used to assess brain activity. Typically, brain activity progresses from: 0 spikes (post-ictal depression); to 1 spike (∼13 min); to 2 spikes (∼18 min); to 3 spikes (21 min); to 4 spikes (24 min); 5 spikes (27 min); and to 6 spikes (29 min). The slope of this seizure progression is a key metric for seizure susceptibility (see **Fig. 3b**). Intermittently during CHS, mice would have a discrete seizure lasting over 30 s; another metric for seizure susceptibility (see **Fig. 3c**). After 30 min the stimulation was stopped, because the mice entered status epilepticus. In this study, a large proportion of mice died after 30 min of status due to tonic phase apnea, as observed in other mouse models of epilepsy (Wenker et al., 2021).

To study the effect of microglia depletion on epilepsy we electrically kindled VGAT-Cre mice after IP injection of 10-15 mg/kg kainic acid (hybrid kindling). Mice were maintained on control chow (low PLX3397) until the mice developed epilepsy, which is defined as having 2 or more spontaneous seizures (King et al., 1998). Mice were then randomized to stay on control chow or switched to high dose PLX3397 chow (N=14). Video/EEG recording were continued for at least 3 weeks to quantify spontaneous recurring seizures (SRS). As noted previously, epilepsy in VGAT-Cre mice spontaneously remits (Straub, Vitko, Gaykema, & Perez-Reyes, 2021).

### 2.5 Open Field Behavioral Test

The open field test was carried out in a custom-built arena using a white plastic material with 35 (L) x 35 (B) x 21 (H) cm dimensions. Mouse cages were moved into the testing room and allowed to acclimate for 1 hour. Illumination in the room was maintained at 150 Lux intensity, temperature and relative humidity were also relatively constant at 70.2 ± 0.9°F and 40.9 ± 5%, respectively. All experiments were done during the light cycle. Four to five mice from the same cage were simultaneously placed into different arenas that had been cleaned with 70% ethanol. Each mouse was placed adjacent to the wall of the arena and allowed to freely explore the space. Their open field activities (horizontal locomotion and mobility) in the arenas were video monitored for 10 min using EthoVision® XT (Noldus, Wagenigen, Netherlands), and then subsequently tracked offline for activity analysis using the same software.

### 2.6 Chronic window implantation and two photon imaging

Mice were implanted with a chronic cranial window as previously described (Bisht, Sharma, & Eyo, 2020). Briefly, during surgery, mice were anesthetized with isoflurane (5% for induction; 1-2% for maintenance) and placed on a heating pad. Mice were treated subcutaneously at the site of surgery with 100μL of 0.25% bupivacaine as a local anesthetic before cutting into the skin above the head for the craniotomy. Using a dental drill, a circular craniotomy of > 3 mm diameter was drilled at 2 mm posterior and 1.5 mm lateral to bregma, the craniotomy center was around the limb/trunk region of the somatosensory cortex. A 70% ethanol-sterilized 3mm glass coverslip was placed inside the craniotomy. A light-curing dental cement (Tetric EvoFlow) was applied and cured with a Kerr Demi Ultra LED Curing Light (DentalHealth Products). iBond Total Etch glue (Heraeus) was applied to the rest of the skull, except for the region with the window. This was also cured with the LED light. The light-curing dental glue was used to attach a custom-made head bar onto the other side of the skull from which the craniotomy was performed. Mice were treated subcutaneously with buprenorphine slow release (SR) immediately after the surgery as an analgesic. Mice were allowed to recover from anesthesia for 10 minutes on a heating pad before returning to their home cage. Mice were allowed to recover from the cranial window surgery for 2-4 weeks before commencement of chronic imaging. Only surviving mice with a clear glass window were used for the imaging studies.

For chronic imaging, mice were anesthetized with isoflurane. The head of the anesthetized mice was stabilized and mounted by the head plate and the animal was placed on a heating plate at ∼35°C under the two-photon microscope. A hundred μl of Rhodamine B dye (2 mg/mL) was injected intraperitoneally to label the vasculature. As previously described, for longitudinal imaging, the blood vessel architecture visible through the craniotomy window was carefully recorded as a precise map of the brain region being visualized and was used to trace back to the original imaging site for chronic imaging studies (Bisht et al., 2020). Imaging was conducted using a Leica SP8 Multiphoton microscope with a coherent laser. A wavelength of 880 nm was optimal for imaging both microglia and the blood vessel dye as well as tdTomato. The power output at the brain was maintained at 25 mW or below. Images were collected at a 1024 × 1024 pixel resolution using a 25X 0.9 NA objective at a 1.5X optical zoom. Several fields of view of z-stack images were collected every 1-2 μm through a volume of tissue and used for analysis. To observe microglial dynamics, z-stack images were acquired every minute at 2 μm steps in depth.

### 2.7 Tissue preparation and immunostaining

For confocal microscopy studies, mice were anesthetized with 5% isoflurane, and transcardially perfused with sodium phosphate buffer (PBS; 50 mM at pH 7.4) followed by 4% paraformaldehyde (PFA). All perfusion solutions were chilled on ice prior to use. We used a perfusion pump (Masterflex ® Ismatec ®) at a perfusion flow rate of 7mL/min. Brains were then fixed in 4% PFA overnight. Using a vibratome (Leica VT100S), 50µm thick sections of the brain were cut in chilled PBS. Slices were then stored in cryoprotectant (40% PBS, 30% ethylene glycol, and 30% glycerol) at -20°C while further processing took place. Brain sections containing the ventral hippocampus CA1 (Bregma −3.27 and −4.03), the frontal cortex (Bregma 2.93 and –2.57), and sensorimotor cortex (Bregma –2.5 and +2.0) were examined. For WFA (1:800, Wako, # 019-19741) immunohistochemical staining for fluorescence microscopy analysis, brain sections were washed in PBS, blocked with blocking buffer, and incubated overnight at 4°C with primary antibody solution against WFA. After washing, brain slices were incubated in appropriate secondary antibody solution for 1 hour at room temperature

### 2.8 Fluorescence Microscopy

Fluorescently immunolabelled brain sections were imaged with a confocal microscope or an EVOS microscope. Image analysis to quantify the WFA area, cell number staining intensity as well as the number of cFos^+^ cells through z-stacks was done using ImageJ. WFA cell number was automatically counted using the ImageJ cell counter plugin. The WFA staining intensity was quantified as a measure of the florescence intensity for WFA and normalized to the area of the interest. The perineuronal net (PNN) area was quantified as a ration of the total area occupied by WFA signal over the total area of the tissue examined. cFos+ cells were identified in z-stack images as distinct from Rhodamine labelled blood vessels in two ways. First, morphological distinctions through the z-stack could be observed as blood vessels are tubular and can be determined through the z-stack and cFos^+^ cells often displayed a cell body with radiating florescent signals. Second, while blood vessels were always present in images from adjacent weeks, cFos+ cells sometimes were absent in a previous week and appeared in a subsequent week.

### 2.9 Tissue processing and flow cytometry

Mice were euthanized with CO_2_ and sprayed with 70% ethanol. The spleen was collected and placed in cold complete RPMI media (cRPMI) (10% FBS, 1% Sodium pyruvate, 1% non-essential amino acids, 1% penicillin/streptomysin, 0.1% 2-ME). The spleen was mashed through a 40-µm filter in 50mL conical using a syringe plunger and through with 15-mL cRPMI. The suspension was centrifuged at 1600 rpm for 5 min at 4°C using Eppendorf Centrifuge 5804R with an S-4-72 rotor. The spleen was resuspended in 2mL RBC lysis buffer for 2 min and the reaction was stopped by adding 13mL cRPMI. The suspension was once again centrifuged, and the resulting pellet were resuspended in 5mL cRPMI and kept on ice. The femur and tibia were carefully removed without splintering them to obtain the bone marrow. The bones were placed in a petri dish containing 70% ethanol for 1-2 minutes to disinfect and transferred to a petri dish containing 4mL cRPMI on ice. The femora and tibiae were flushed with 10mL ice cold cRPMI using a 21G needle attached to a 10mL syringe. A single-cell suspension was generated by gently triturating the cells through the needle until large clumps were no longer present and the suspension was ran through a 40µm filter in 50mL conical tube. The suspension was centrifuge at 1600 rpm for 5 min at 4°C. The bone marrow was resuspended in 2mL RBC lysis buffer for 2 min and the reaction was stopped by adding 13mL cRPMI. The suspension was centrifuged at 1600 rpm for 5 min at 4°C, the supernatant was aspirated and the pellet was resuspended in 5mL cRPMI. Following the generation of a single-cell suspension, cells were counted using automated cell counter (C100, RWD, China), 150 µL (∼1×10E6) of each sample were placed in a 96-well plate and incubated for 10 minutes in 50 µL Fc block (1:1000, CD16/32, Clone 93, eBioscience) at room temperature. Cells were then incubated in primary antibodies at a concentration of 1:200 and fixable viability dye eFluor 506 (eBioscience) at a concentration of 1:800 for 30 minutes at 4°C. Antibody clones used for experiments included: NK1.1 (PK136), CD3e (145-2C11), CD8a (53-6.7), CD4 (RM4-5), CD19 (eBio 1D3), Ly6C (HK1.4), F4/80 (BM8), CD11b (M1/70), Ly6G (1A8), CD45 (30-F11), MHC II (M5/114.15.2), and CD11c (N418) (Invitrogen). After staining, cells were washed twice and fixed overnight in 2% PFA at 4°C. Cells were washed twice with cell staining buffer (BioLegend, Cat. # 420201) and transferred into a 5mL Polystyrene round-bottom tube with cell-strainer cap tubes (Falcon), then were analyzed on a Gallios flow cytometer (Beckman-Coulter) by gating on 250,000 live events. Flow cytometry data was analyzed using FlowJo (version 10.8.1).

### 2.10 Statistical analysis

Data were initially measured for normality and homoscedasticity and upon comparing normal distributions and variances further analyzed with the respective tests. Student’s t-test was used to compare two-groups. Other comparisons were evaluated using one-way ANOVA (more than 2 groups) or two-way ANOVA (experiments with 2 variables), followed by suitable post hoc test for multiple comparisons within the tested groups. Other specific tests are stated in the figure legend for each of the experiments.

## 3. RESULTS

### 3.1 Pharmacogenetic microglial elimination worsens kainic acid-induced seizures

To begin, we attempted to confirm previous results showing that pharmacogenetic microglial elimination worsens chemoconvulsive seizures (Mirrione et al., 2010; Wu et al., 2020) by using a pharmacogenetic model of microglial elimination in a chemoconvulsive seizure model. We used IP injection of the chemoconvulsant, kainic acid (KA), a glutamate receptor analog at 24-27 mg/kg to investigate seizure severity (**Supplemental Figure 1**). CX3CR1^Cre^ mice were crossed with Rosa26^iDTR^ mice to generate CX3CR1^Cre/+^:Rosa26^iDTR/+^ that express the diphtheria toxin receptor (DTR) on CX3CR1^+^ cells (**Supplemental Figure 1a**). In adulthood, Rosa26^iDTR/+^ mice treated with saline or diphtheria toxin (DT, 50 µg/kg) and therefore not expected to have depleted microglia showed similar seizure scores when injected with KA a day after DT treatment (**Supplemental Figure 1b-c**) indicating that DT treatment in microglial-sufficient Rosa26^iDTR/+^ mice does not alter seizure severity. Interestingly, compared to saline-treated CX3CR1^Cre/+^:Rosa26^iDTR/+^ mice which are microglia-sufficient, DT-treated CX3CR1^Cre/+^:Rosa26^iDTR/+^ mice which become microglia depleted exhibited more severe seizures (**Supplemental Figure 1d-e**) and all the mice died shortly after an hour of KA treatment (**Supplemental Figure 1d**) indicating aggravated seizures with pharmacogenetic microglial elimination.

### 3.2 Pharmacological microglial elimination and repopulation

The above results are consistent with previous pharmacogenetic studies that eliminated microglia in chemoconvulsive seizures (Wu et al., 2020). However, recent findings suggest that pharmacogenetic approaches to eliminate microglia result in additional confounds ranging from altered brain ventricles as a structural abnormality, to increased neuronal cell death, to ataxic behavior, and to increased glial reactivity and cytokine levels (Bedolla et al., 2022; Rubino et al., 2018). Therefore, to clarify microglial contributions to seizures, we employed a pharmacological microglial elimination approach using treatment with PLX3397 (PLX), a CSF1R antagonist. CSF1R is required for microglial survival. Therefore, PLX treatment efficiently eliminates microglia without inflammatory reactions or altered brain structures (Bedolla et al., 2022; M. R. Elmore et al., 2014; Green, Crapser, & Hohsfield, 2020; Szalay et al., 2016). We confirmed that PLX (660mg/kg) delivered through the chow progressively eliminated microglia by 7 days (**Figure 1a-b**). As with PLX-treated mice that we refer to as “microglial depleted” mice, “control” mice received chow with either 0 mg/kg PLX or 75 mg/kg for 7 days. The latter concentration was previously shown to have minimal brain penetration of PLX, but significant plasma concentration that was higher than the concentration required in the brain to eliminate microglia (M. R. Elmore et al., 2014). We confirmed that the 75mg/kg PLX dose did not affect microglial density (**Figure 1g**) compared to the 660mg/kg dose (**Figure 1b**). These “control” mice we refer to as “microglia sufficient” mice. Interestingly, although PLX5622 was recently shown to reduce lymphoid and myeloid cell numbers with a 3-week treatment (Lei et al., 2020; Spiteri et al., 2022), we speculated that the prolonged treatment regimen could have yielded this result. Therefore, we used PLX at a high concentration (660mg/kg) to facilitate rapid microglial cell elimination. We found that a one-week PLX3397 treatment did not elicit significant changes to immune cell numbers in the spleen or bone marrow except for a slight but significant increase in monocyte cell numbers (**Figure 1c-d**). To assess whether microglia repopulate the brain rapidly following PLX3397 withdrawal, treated mice with either control or PLX chow for a week, returned all mice to control chow for another week and noted that microglia had repopulated the brain by 7 days (**Figure 1e-f**) Therefore, we used this approach of pharmacological microglial elimination and repopulation to interrogate microglial roles in seizure severity.

**FIGURE 1.**
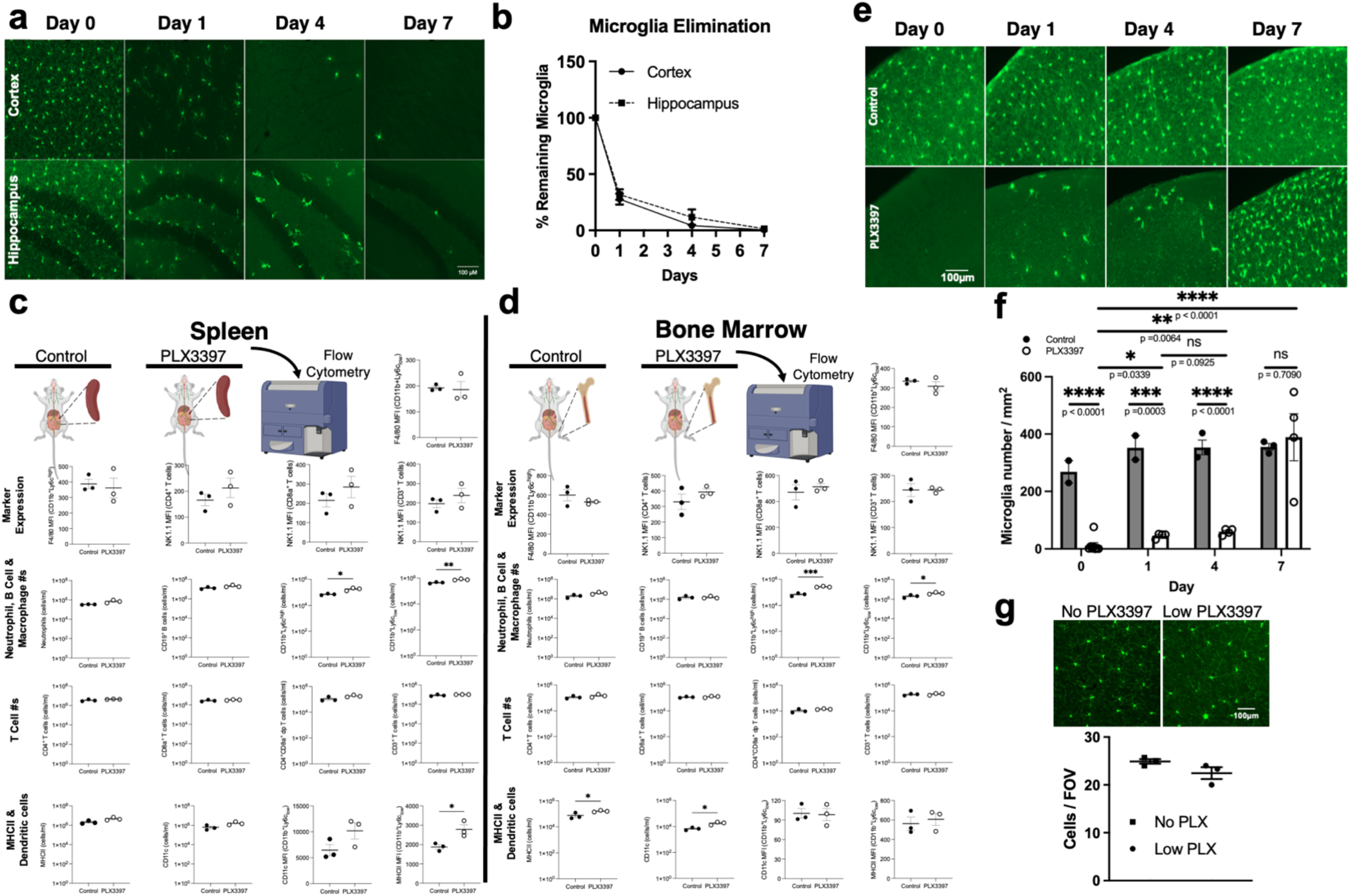
Pharmacological microglial elimination and repopulation. a-b, Representative images (a) and quantification of the percent of microglial cells (b) from the cortex and hippocampal dentate gyrus of CX3CR1^GFP/+^ treated with PLX3397 chow (660mg/kg) at 0, 1, 4 and 7 days of treatment. n = 3 mice per group. **c-d**, Schematic representation of splenic (c) and bone marrow (d) isolation following a 7-day treatment with control of PLX3397 chow for processing by flow cytometry of various immune cells including T cells, B cells, neutrophils, inflammatory and patrolling monocytes. n = 3 mice per group (e-g), Representative images (e) and quantification (f-g) of the density of microglial cells from the cortex of CX3CR1^GFP/+^ treated with control or PLX3397chow for 7 days and then withdrawn from the PLX3397 chow at 0, 1, 4 and 7 days. n = 2–4 mice per group. Data presented as mean ± s.e.m f and g. Statistics calculated by Student’s T-test, *p < 0.05; **p < 0.01; ***p < 0.001, ****p < 0.0001.

### 3.3 Pharmacological microglial elimination worsens kainic acid-induced seizures

Mice were treated with either control or PLX chow for 7 days before KA treatment (**Figure 2a**). PLX-treated (and therefore microglial deficient) mice showed worsened status epilepticus to KA (**Figure 2b-c**) when compared to control-treated (and therefore microglial sufficient) mice without altering mortality (**Figure 2d**). Seizure severity differences began to be apparent as early as 30 mins into KA treatment and persisted through 4 h (**Figure 2b; Supplemental Figure 2a-e**). Furthermore, increased seizure severity in microglial-deficient mice compared to microglial-sufficient mice was observed in both male and females (**Supplemental Figure S2f-i**).

**FIGURE 2.**
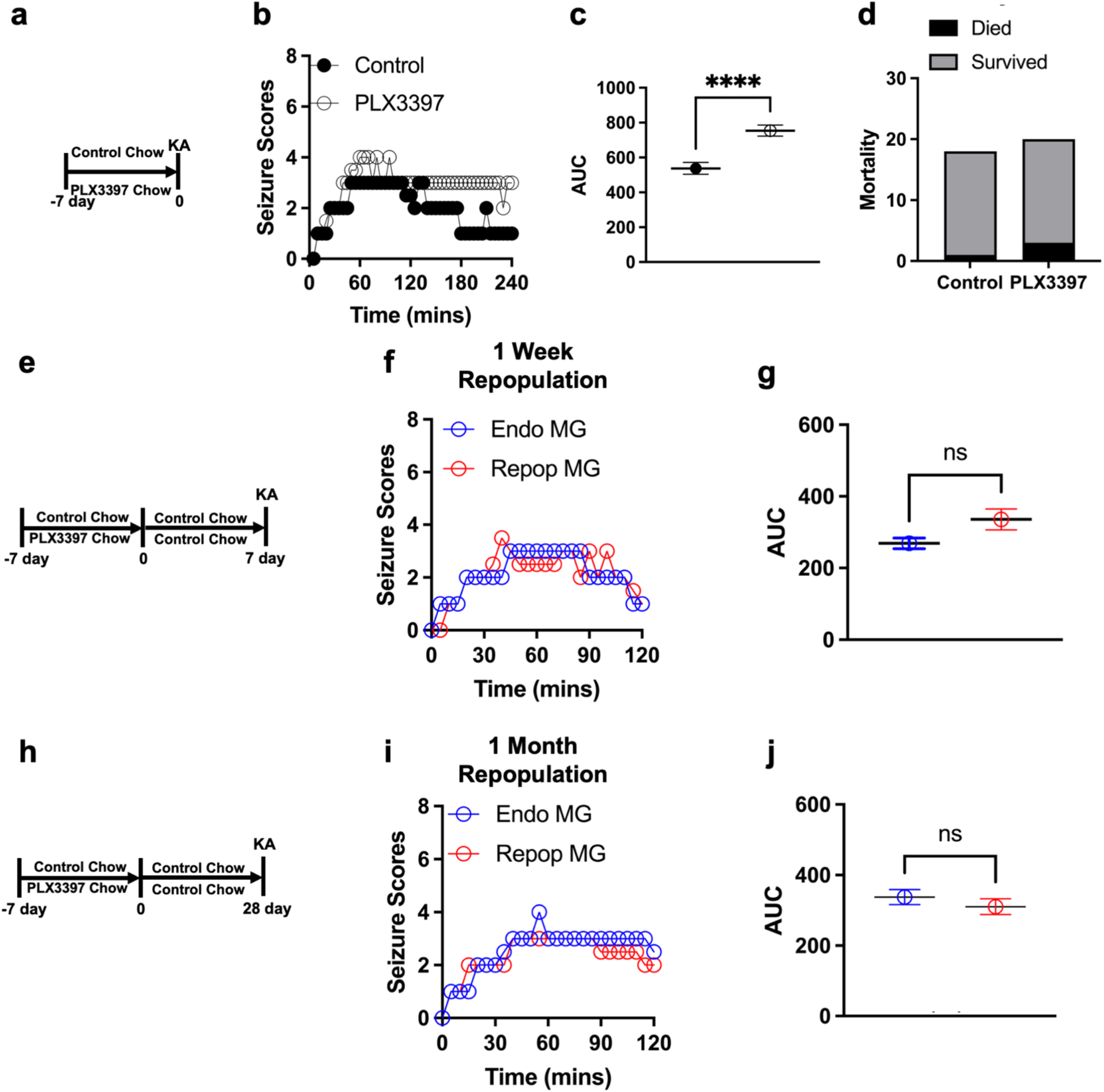
Endogenous and repopulated microglia are protective in chemoconvulsive seizures. (a) Diagram of experimental scheme for microglial depletion and chemoconvulsive kainic acid (KA) seizures. (b and c) Overall (b) and area under the curve (AUC, c) Racine seizure scores from control and PLX3397 treated mice monitored over 240 mins (4h). (d) Percent animal survival by 4h of KA treatment. n = 18-20 mice per group. (e) Diagram of experimental scheme for microglial depletion and subsequent repopulation followed by KA seizure induction after 7 days. (f and g) Overall (f) and AUC (g) Racine seizure scores from control and PLX3397 treated mice monitored over 120 mins (2h). n = 20 mice per group. (h) Diagram of experimental scheme for microglial depletion and subsequent repopulation followed by KA seizure induction after 28 days. (i and j) Overall (i) and AUC (j) Racine seizure scores from control and PLX3397 treated mice monitored over 120 mins (2h). n = 16-20 mice per group. Data presented as median in b, f and I and mean ± s.e.m in c, g and j. Statistics calculated by Student’s T-test and Chi-squared with Fisher’s exact test in d. ****p < 0.0001.

**FIGURE 3.**
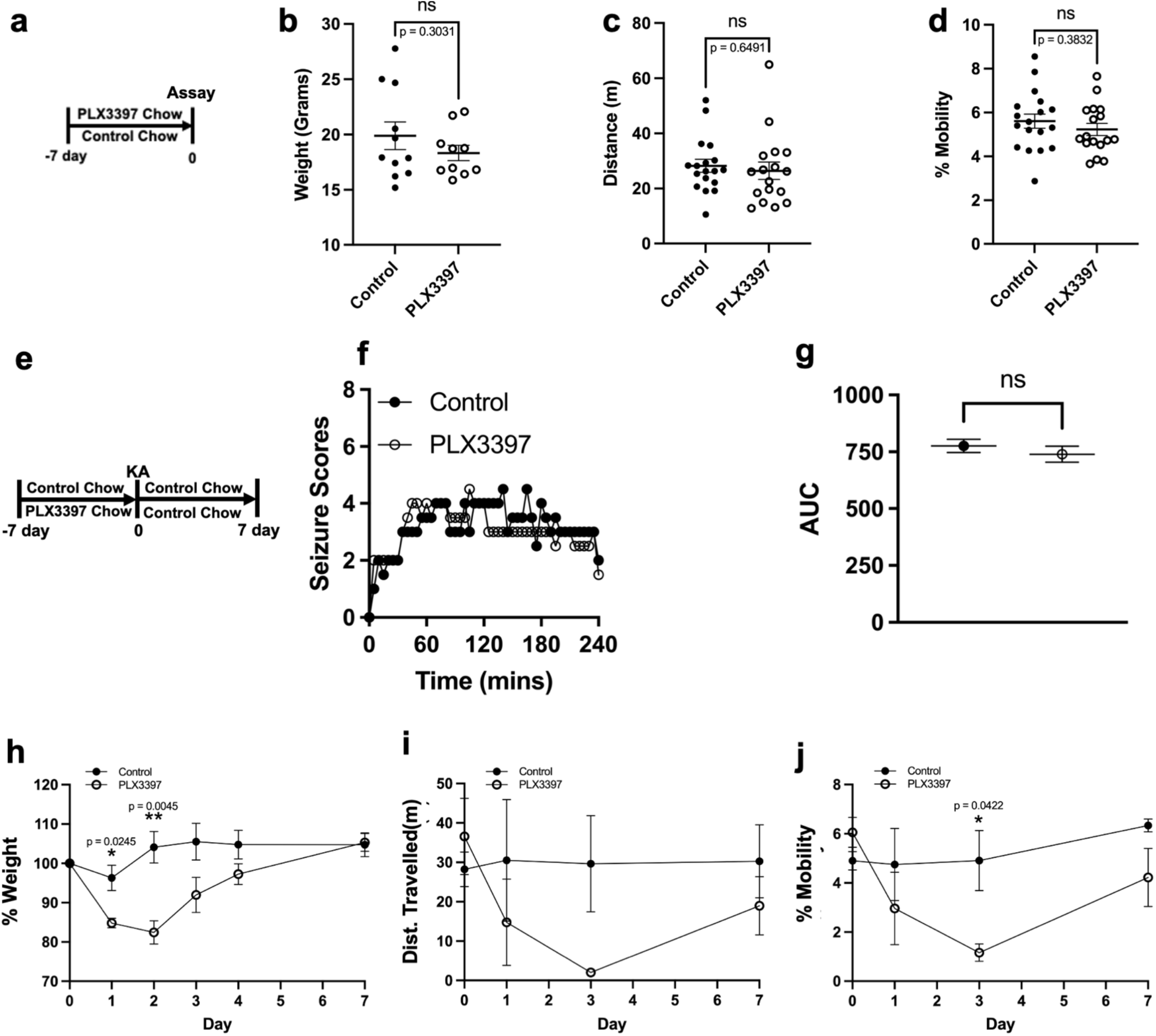
Microglia are important in the recovery from chemoconvulsive seizures. (a) Diagram of experimental scheme for microglial depletion and behavioral assays. (b-d) Mice weights (b) distance travelled (c) and degree of mobility (d) exhibited by mice in an open field without KA treatment. n = 10-18 mice per group. (e) Diagram of experimental scheme for microglial depletion followed by KA seizure induction after 7 days and behavioral assays for 7 days. (f and g) Overall (f) and area under the curve (AUC, g) Racine seizure scores from control and PLX3397 treated mice monitored over 240 mins (4h). n = 8 mice per group. (h-j) Mice weights (h) distance travelled (i) and degree of mobility (j) exhibited by mice in an open field following KA treatment. n = 4-6 mice per group. Data presented as mean ± s.e.m except in f where data is presented as median. Statistics calculated by Student’s T-test for b, c, d and g and Two-way ANOVA for h-j. *p < 0.05; **p < 0.01.

Next, we examined the level of cell activity following KA treatment in the presence or absence of microglia. To this end, we used TRAP2 (targeting recombination in active populations) mice crossed with floxed tdTomato reporter mice (DeNardo et al., 2019; Madisen et al., 2010) where tamoxifen (TAM) treatment induces tdTomato expression under the control of the cFos promoter. cFos is an immediate early gene expressed by active cells. Littermate TRAP2 mice were treated with either PLX or control chow for 7 days to eliminate microglia. After this, mice were treated with KA and at 3 h, mice were injected with TAM (40 mg/kg) and then euthanized after 7 days to allow for tdTomato expression (**Supplemental Figure S3a**). With KA treatment, PLX-treated (and therefore microglial deficient) mice displayed increased cFos expression in both the cortex and hippocampus compared to control (and therefore microglial sufficient) mice (**Supplemental Figure S3b**).

To further visualize these changes *in vivo*, we crossed the TRAP2 mice with the microglial GFP reporter mice to generate double transgenic “TRAP-GFP” mice and conducted longitudinal *in vivo* two photon imaging (**Supplemental Figure S3c**). Consistent with the above findings in TRAP2 mice, longitudinal imaging of the same fields of view in TRAP-GFP mice revealed that KA treatment induced the expression of cFos in new cells (arrows in **Supplemental Figure S3d**, quantification in **S3e**). Interestingly, microglial elimination with PLX did not itself increase cFos expressing cell numbeers but coupled with subsequent KA treatment resulted in a noticeable labelling on several cortical cells (arrows in **Supplementary Figure S3d**, quantification in **S3e**).

Because inflammatory mediators are correlated with seizure severity (Choi, Min, & Shin, 2011; Terrone, Frigerio, Balosso, Ravizza, & Vezzani, 2019; Wang & Chen, 2018), we also evaluated inflammatory cytokine expression following KA-induced seizures in control (and therefore microglial sufficient) and PLX-treated (and therefore microglial deficient) littermate mice. We examined tissue expression of various cytokines at 3 h of KA treatment when cytokines were previously shown to be significantly elevated in this model of seizures (Avignone et al., 2008). We noted no differences between the microglial deficient and microglial sufficient groups (**Supplemental Figure S4**) suggesting that cytokine levels following KA treatment are maintained independent of microglia.

In addition, perineuronal nets (PNNs) are components of the extracellular matrix whose modulation can regulate seizure phenotypes (Chaunsali, Tewari, & Sontheimer, 2021). Since microglia regulate the ECM, it is therefore possible that they may influence seizure phenotypes by modulating PNNs. However, consistent with a previous report that documented a lack of gross alterations in PNN density following a one-month PLX treatment (Strackeljan et al., 2021), our PLX treatment did not alter gross perineuronal net (PNN) density or intensity in either adult (**Supplemental Figure S5a-d**) or neonatal mice (**Supplemental Figure S5e-h**) indicating that the increased KA-induced seizures in with microglial elimination occurs independent of tissue PNN density in the KA model of seizures. Together, these results indicate that a neither a cytokine-dependent or a PNN-dependent mechanism is sufficient to explain the exacerbated KA-induced seizures with microglial elimination.

### 3.4 Endogenous and repopulated microglia are similarly beneficial in acute kainic acid-induced seizures

Eliminating and repopulating microglia following PLX-induced elimination is neuro-beneficial with aging (M. R. P. Elmore et al., 2018) and following intracerebral hemorrhage (X. Li et al., 2022). Therefore, we speculated that repopulated microglia may have a stronger seizure-limiting potential than endogenous microglia. To this end we treated mice with either control or PLX chow for a week, returned all mice to control chow for another week by which time microglia had repopulated the brain (**Figure 1e-f**) and then exposed all mice to KA (**Figure 2e**). Although both cohorts of mice were microglial sufficient, we referred to the mice previously exposed to PLX as the “repopulated microglia” [Repop MG] and the mice treated with control chow as the “endogenous microglia” [Endo MG] group. Seizure severity was similar in mice with endogenous and repopulated microglia (**Figure 2f-g**). Because microglia are known to progressively mature during repopulation following PLX treatment (Zhan et al., 2019), we asked whether more mature repopulated microglia could show a more protective effect by conducting KA-induced seizures 4 weeks after PLX treatment when microglial maturation following PLX is complete (Zhan et al., 2019). As with one-week, mice with repopulated microglia at one-month showed similar seizure phenotypes to mice with endogenous microglia during KA seizures (**Figure 2h-j**). Together, these results indicate that both endogenous and repopulated microglia play seizure-limiting roles in chemoconvulsive seizures.

### 3.5 Microglia play beneficial roles in the recovery from kainic acid-induced seizures

Next, we attempted to determine microglial contributions to recovery from seizures. At 7 days of PLX or control treatment, we measured mice weight and open field behaviors. By itself, PLX did not affect mouse weight, distance travelled, or mobile activity in an open field (**Figure 3a-d**). Therefore, following a 7-day PLX or control treatment, both groups of mice were treated with KA and returned to control chow. We selected mice that exhibited similar seizure severity whether previously treated with control or PLX chow and monitored them for 7 days. Mouse weight and activity in an open field were monitored for 7 days (**Figure 3e**). For these studies, we selected only mice that showed similar seizure scores as they stayed or were in and out at least seizure stage 4 between the 1^st^ and 3^rd^ hours of seizures independent of PLX treatment (**Figure 4f-g**) to account for a similar seizure severity. Nevertheless, PLX-treated mice displayed increased weight reductions (**Figure 4h**), reduced distance travelled (**Figure 4i**), and reduced mobile activity (**Figure 4j**, see **Supplemental Video S1- 3**) in an open field during the earlier phases of recovery when microglia were still reduced following PLX withdrawal (**Figure 2e-g**). These deficits were restored by 7 days (**Figure 4f-h**) when microglia had repopulated the brain (**Figure 2e-g**) suggesting a correlation between microglial absence/reduction and behavioral deficits on the one hand and microglial repopulation and behavioral recovery on the other. Together, these results indicate that microglia are important for containing seizures and facilitating recovery from seizures.

**FIGURE 4.**
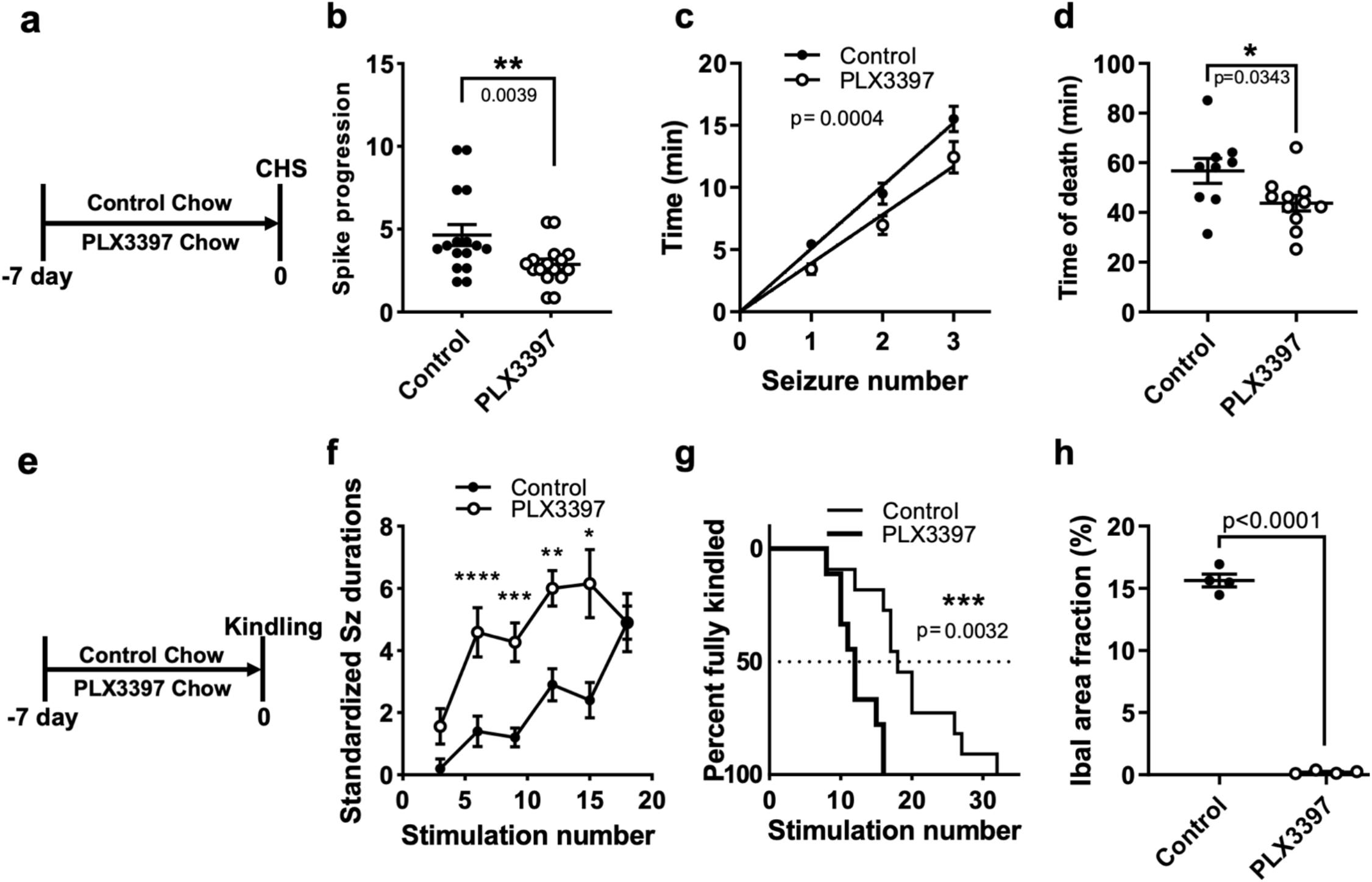
Microglia are beneficial in electrically induced seizures. (a) Diagram of experimental scheme for microglial depletion and chronic hippocampal stimulation (CHS) after 7 days using C57Bl/6J mice (n=16 per group). (b) Evolution of spikes during CHS stimulation (see Methods for details). (c) Time to CHS-evoked discrete seizures. (d) Time to CHS-induced death from start of CHS. (e) Diagram of experimental scheme for microglial depletion for 7 days followed by electrical kindling. (f) Standardized seizure durations elicited during kindling of VGAT-Cre mice. n = 9 – 11 mice per group. (g) Kaplan-Meyer plot of the number of stimulations required to elicit the full kindled state (5 evoked bilateral tonic-clonic seizures with loss of balance) in VGAT-Cre mice. (h) Quantification of Iba1^+^ cells in control and PLX3397-treated mice. n = 4 mice per group. Data presented as mean ± s.e.m. Statistics calculated by Student’s t-test (b, d, f), linear regression (c), Log-rank Mantel Cox test (g), or 2-way ANOVA followed by Sidak’s test (h). Significance is denoted by: *, p < 0.05; **, p < 0.01; ***, p < 0.001 and ****, p < 0.0001.

### 3.6 Pharmacological microglial elimination worsens seizure phenotypes in electrically-induced seizure paradigms

Next, we sought to extend our findings in the chemoconvulsive paradigm to other seizure paradigms. Therefore, we employed a continuous hippocampal stimulation (CHS) model (Lewczuk et al., 2018). Following implantation of EEG electrodes, we treated mice with PLX or control chow for 7 days and then performed CHS (**Figure 5a**). Here, microglial elimination accelerated the progression to status epilepticus (**Figure 5b**), accelerated the appearance of discrete seizures during CHS (**Figure 5c**), and accelerated animal mortality (**Figure 5d**). Furthermore, we used our recently developed VGAT-Cre model of temporal lobe epilepsy (TLE) where VGAT-Cre mice develop spontaneous seizures following mild electrical stimulation to induce kindling (Straub et al., 2020). As with CHS, following implantation of EEG electrodes, mice were treated with PLX or control chow for 7 days and a kindling regimen was implemented, which consisted of kindling six times a day with each kindling session separated by at least an hour (see Methods for details; **Figure 5d**). PLX-treated VCAT-Cre mice experienced a longer duration of evoked seizures (**Figure 5f**), and displayed an acceleration to fully the kindled state (5 evoked convulsive seizures, BSS≥5; **Figure 5g**). Moreover, we confirmed that PLX-treatment indeed eliminated microglia in this model (**Figure 5h**). Together, these results indicate that microglial elimination facilitates the development of electrically induced seizures.

**FIGURE 5.**
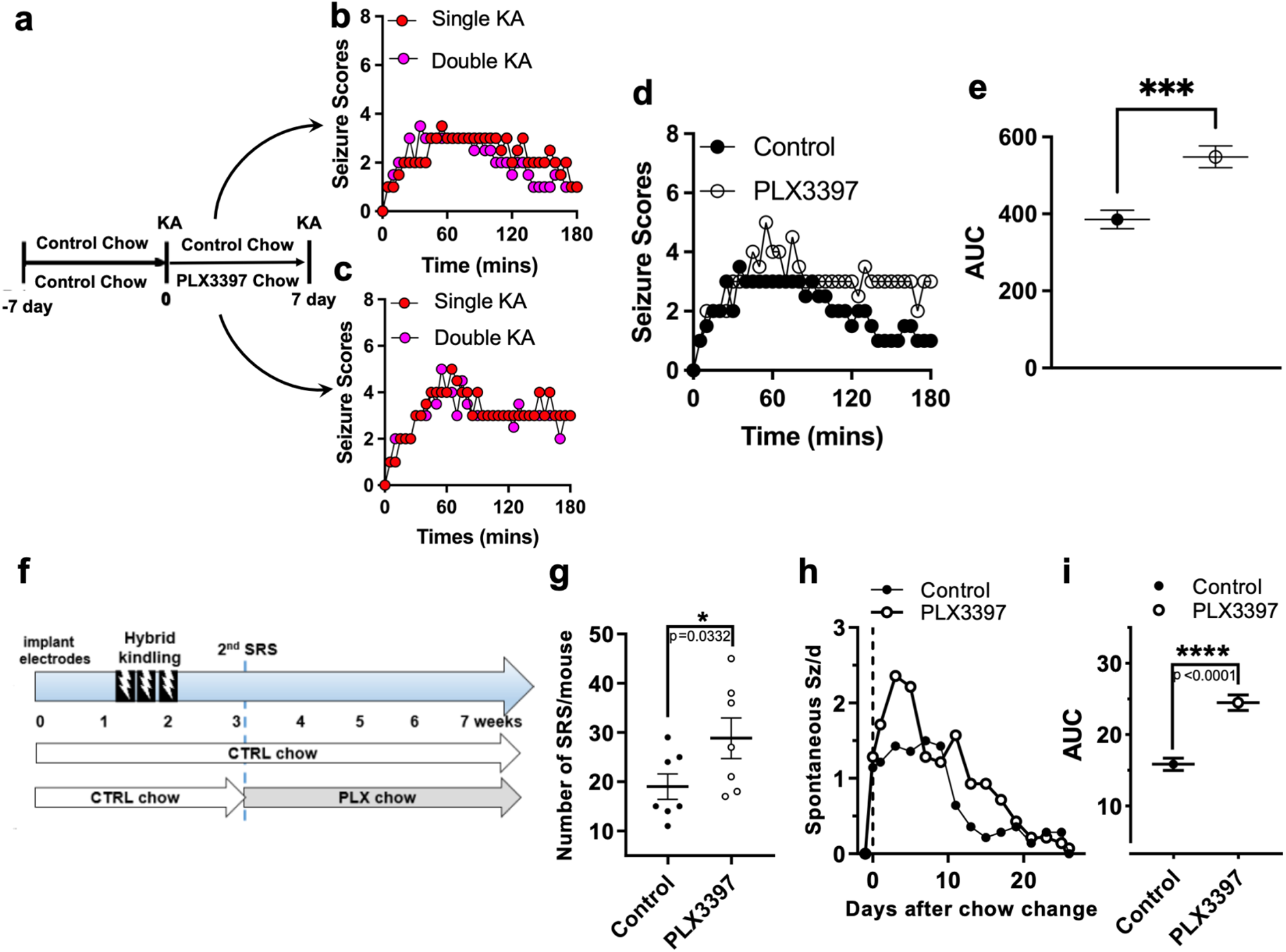
Microglia are beneficial following secondary and spontaneously occurring seizures. (a) Diagram of experimental scheme for two KA exposures with a 7-day microglial depletion between the KA treatments. (b and c) Median Racine seizure scores from a single and following a double bout of KA treatment in control (b) and PLX3397 (c) treated mice n = 10-20 mice per group. (d and e) Overall (d) and average (e) Racine seizure scores from a second bout of KA-induced seizures following control and PLX3397 treatment monitored over 180 mins (3h). n = 8 mice per group. (f) Diagram of experimental scheme for hybrid kindling followed by monitoring of spontaneous recurrent seizures (SRS) with microglial depletion. (g) Quantification of the number SRS over a 2-week period with microglial depletion. (h and i) Time course of spontaneous seizures after chow was changed (day 0, binned in 2 day intervals). n = 7 mice per group Data were analyzed by area under curve (AUC) and the results are plotted in (i). Data presented as median in b – d and mean ± s.e.m in e, g and i. Statistics calculated by Student’s t-test *p < 0.05; ****p < 0.0001.

### 3.7 Pharmacological microglial elimination worsens seizure phenotypes in subsequent induced or spontaneous seizures

Given the evidence above of microglia playing beneficial roles in primary or initial seizures, we sought to investigate possible roles for microglia in secondary or subsequent seizures to an initial seizure. We interrogated this possibility first using our chemoconvulsive model. Two groups of mice were treated with control chow for 7 days and exposed to KA. One group was then maintained in control chow and the second group in PLX chow to deplete their microglia. After another 7 days of the respective chow treatment, mice were treated with a second round of KA (**Figure 6a**) and their seizure scores monitored over time. Interestingly, seizures scores were similar between microglia-sufficient mice that experienced a single or two rounds of KA-induced seizures (**Figure 6b**) as well as between microglia-deficient mice that experienced a single or two rounds of seizures (**Figure 6c**) indicating that an initial KA-induced bout of seizures does not potentiate a second KA-induced bout of seizures 7 days later. However, in the second KA-induced bout of seizures experience, mice that lacked microglia between the first and second bouts also showed a greater seizure severity compared to mice that were microglia-sufficient (**Figure 6d-e**).

**FIGURE 6.**
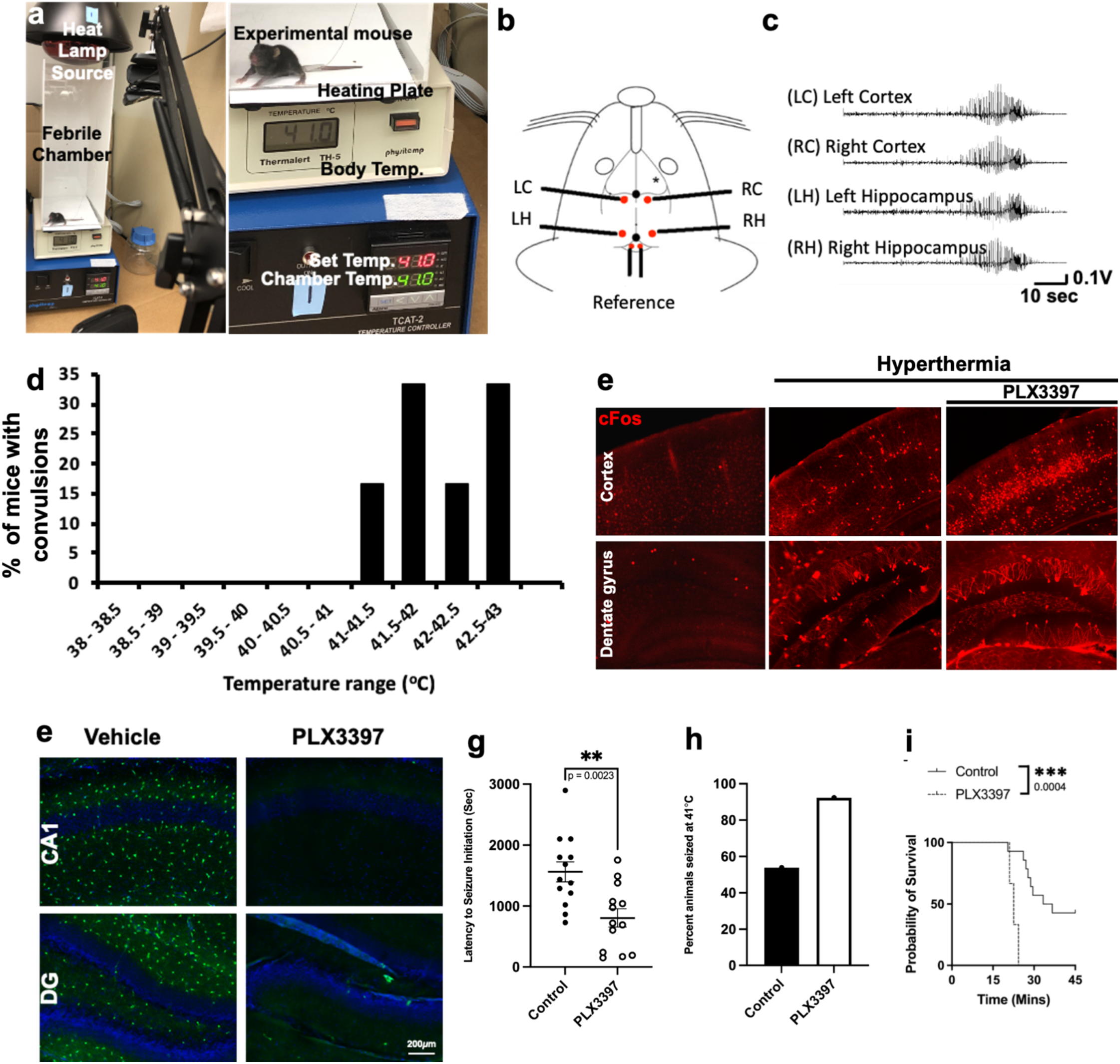
Microglia are beneficial in hyperthermia-induced seizures in developing mice. (a) Image of hyperthermic (febrile) seizure setup with a mouse in a chamber made of plexiglass with heat provided by a heating plate below and a heat lamp from above. Temperature in chamber can be set and measured externally. (b) Schematic of a mouse head with implanted electrode in the left cortex (LC), right cortex (RC), left hippocampus (LH) and right hippocampus (RH). (c) Electrical activity is detected in all four regions upon hyperthermia exposure. (d) The percent of mice that show behavioral convulsions under hyperthermia at the different temperatures. (e) cFos expression in the cortex and dentate gyrus of normothermic and hyperthermic conditions as well as with microglial depletion. (f) Efficient microglial elimination with PLX3397 treatment in developing mice. (g-i) Microglial elimination and its effects of time to the first seizures (g) percent of mice that seize at 41°C (h) and mortality rate (i) during hyperthermia-induced seizures in developing mice. n = 13 mice per group. Statistics calculated by Student’s T-test and Chi-squared with Fisher’s exact test in i. **p < 0.01; ****p < 0.0001.

Furthermore, we extended these studies into a paradigm where electrically induced seizures can generate spontaneous recurrent seizures (SRS) in VGAT-Cre mice. To this end, electrode-implanted mice were kindled using a hybrid kindling paradigm (see Methods) and then allowed to recover and develop spontaneous seizures over a 2-week period while SRS were measured (**Figure 6f**). Consistent with our findings with KA, PLX-treated mice showed an increase in the number of SRS in the chronic phase (**Figure 6g**). These results in both a chemoconvulsive and electrical seizure model indicate that that microglia also play beneficial roles in subsequent/secondary seizures following an initial seizure.

### 3.8 Pharmacological microglial elimination worsens seizure phenotypes in hyperthermia-induced seizures

Finally, we sought to translate these findings to a more clinically relevant seizure paradigm. Febrile seizures (FS) are the most common neurological disorder in children globally. They occur in 2-5% of children in the United States and Western world (Leung, Hon, & Leung, 2018), 4% of children in Tanzania (Winkler, Tluway, & Schmutzhard, 2013) and 7-10% of children in Asia (Byeon, Kim, & Eun, 2018; Hackett, Hackett, & Bhakta, 1997; Tsuboi, 1984). Furthermore, a subset (up to 18%) of these are prolonged and can have long-term effects on children including developmental delays (Hesdorffer et al., 2011; Martinos et al., 2019), increased association with neurodevelopmental disorders (Bertelsen, Larsen, Petersen, Christensen, & Dalsgaard, 2016; Gillberg, Lundstrom, Fernell, Nilsson, & Neville, 2017; Salehi, Yousefichaijan, Safi Arian, Ebrahimi, & Naziri, 2016), and the development of temporal lobe epilepsy (TLE), the most common form of focal epilepsy in adults (Cendes et al., 1993; Lewis et al., 2014; Shinnar, 2003; Yokoi et al., 2019).

Therefore, we developed a febrile seizure system where exposure of postnatal day (P)13-15 mice to increased temperatures in a hyperthermia (HT) chamber (**Figure 7a**) reliably elicited behavioral convulsions at temperatures ≥ 41°C (**Supplemental Video S4**). These convulsions correlated with electrographical seizure activity in both the cortex and hippocampus (**Figure 7b-c**) beginning to occur at 41°C (**Figure 7d**). Importantly, hyperthermia increased cFos expression which was further increased by PLX-induced microglial elimination (**Figure 7e**). Expectedly, compared to vehicle-treated mice, PLX-treated developing mice showed eliminated microglia (**Figure 7f**) and Consistent with findings in other seizure models, microglial-depleted developing mice displayed convulsions earlier (**Figure 7g**) and a greater percent of mice showed convulsions at 41°C (**Figure 7h**) and greater mortality (**Figure 7i**) from the hyperthermic exposure indicating exacerbated hyperthermia-induced seizures in the absence of microglia. Taken together, our results from a chemoconvulsive, two electrical, and a hyperthermia-induced seizures models with non-inflammatory microglial elimination comprehensively suggest beneficial roles for microglia in acute seizures.

## DISCUSSION

The precise role of microglia in seizure disorders has been elusive with suggested beneficial (Liu et al., 2020; Mirrione et al., 2010; Waltl et al., 2018; Wan et al., 2020; Wu et al., 2020; Zhao et al., 2020) and detrimental roles (Abraham et al., 2012; Di Nunzio et al., 2021; Eun et al., 2015; Kim et al., 2015). Using an approach that lacks inflammatory sequelae, glial reactivity, or brain structural aberrations in three different seizure paradigms, we provide comprehensive evidence for beneficial roles for microglia in mitigating seizures. Specifically, we show that: (1) endogenous or repopulated microglia are important in limiting chemoconvulsive, electrical, and hyperthermic seizures; (2) endogenous microglia facilitate recovery from chemoconvulsive seizures and (3) following an initial seizure, endogenous microglia still play seizure-limiting roles in both chemoconvulsive and electrically induced seizures.

Our studies suggest that microglia are important in regulating seizure severity and indicate that microglia are not required for *seizure initiation* as seizures occur in all models tested in the presence or absence of microglia. This is reasonable since neuronal excitability and/or excitation/inhibition imbalance underly seizure initiation. However, our results suggest that microglia play roles that likely facilitate *seizure reduction*. For example, in the chemoconvulsive model where we measured seizure severity temporally, microglial-sufficient and microglial-deficient mice show similar rises in seizure severity during the first hour. However, microglial-deficient mice show a slower decline from peak seizure severity when compared to microglial-sufficient mice. However, since our studies in the chemoconvulsive studies only measured the behavioral convulsions and not the electrical activity, this hypothesis remains to be fully validated.

Our findings are consistent with recent observations that microglia dampen seizure and network excitability (Cserep et al., 2020; Kato et al., 2016; Y. Li et al., 2012; Szalay et al., 2016) through Gi-dependent mechanisms such as microglial surveillance (Merlini et al., 2021) and well as previous results showing that microglia may facilitate the restoration of dysfunctional neuronal structures following seizures (Eyo et al., 2021). They are also consistent with other findings suggesting that microglial elimination enhances pilocarpine and pentylenetetrazol (PTZ) induced seizures and neurodegeneration (Liu et al., 2020) as well as neurodegeneration following kainic acid-induced seizures (Araki, Ikegaya, & Koyama, 2019). However, these previous studies assessed microglial roles usually in one paradigm i.e., chemoconvulsive models. Ours is therefore a first to comprehensively interrogate microglial roles in different seizure paradigms: chemoconvulsive, electrical, and hyperthermic.

Of interest, in our studies using both chemoconvulsive and electrical seizure paradigms, microglia were also protective in subsequent seizures either induced experimentally or spontaneously following kindling (**Figure 6**). This suggests that even in the early stages following an initial bout of seizures microglia provide beneficial roles. Since the mechanisms of seizure induction in chemoconvulsive, electrical, and hyperthermic paradigms differ but microglia are beneficial in all models, it is possible that microglia employ similar beneficial mechanism(s) that once understood could be enhanced to facilitate seizure containment and mitigation of its detrimental consequences. However, it is also possible that microglia employ different mechanisms for seizure mitigation in the different models of seizures. Therefore, while we could not find a cytokine-dependent mechanism in the chemoconvulsive model, we cannot rule out a cytokine-mediated mechanism in either the hyperthermia-induced or electrical models.

While our results provide supportive evidence for a beneficial role for microglia in several acute seizure models, the precise molecular mechanism(s) microglia employ in this context remain to be determined. Recently, microglial Gi-dependent signaling was shown to regulate seizure phenotypes through microglial surveillance (Merlini et al., 2021). Among the many Gi-coupled receptors expressed by microglia, the P2RY12 and CX3CR1 receptors have been shown to regulate injury-induced microglial surveillance (Haynes et al., 2006; Liang et al., 2009). In addition, the findings from the current study are consistent with molecular results that show that microglial P2RY12 play beneficial roles in adult chemoconvulsive (Eyo et al., 2014) and developing febrile (Wan et al., 2020) seizures. Similarly, CX3CR1-depedent mechanisms were shown to be protective beneficial in chemoconvulsive seizures (Eyo, Peng, et al., 2017). Therefore, one possible mechanism of microglial neuroprotection might be via Gi-dependent P2RY12 and/or CX3CR1 regulated seizures. The precise cellular mechanisms by which microglia dampen seizures remain undetermined. Possible mechanisms could include the clearance of excess neurotransmitters, the facilitation of neurotransmitter receptor recycling, the displacement of synapses, the secretion of anti-excitable or pro-inhibitory factors all to reduce neuronal excitability. Future studies will need to test each of these possibilities.

Importantly, it should be noted that our models are not models of epilepsy proper which takes weeks to months to develop and microglial roles in such paradigms could be different from that delineated herein. In addition, given recent evidence for microglial heterogeneity (Masuda, Sankowski, Staszewski, & Prinz, 2020), our approach to broadly eliminate microglia cannot distinguish possibly different roles for distinct subsets of microglia. Indeed, a recent study reports that selectively eliminating proliferating microglia reduced hippocampal neurodegeneration early following status epilepticus and spontaneous seizures during established epilepsy(Di Nunzio et al., 2021). These results therefore suggest that proliferating microglia are enhance neurodegeneration and reduce seizures in epilepsy. Since our approach eliminated all microglia, we cannot rule out this possibility and future more selective manipulations may need to be employed to assess microglial contributions in such contexts.

## AUTHOR COTRIBUTIONS

SG, JB, EP-R and UBE designed the experiments for the project. SG, JB, RPG, JS, KAO, JOU, DHL, BR, MNC, BP and ABC-S conducted experiments for the project. SG, EP-R and UBE wrote the manuscript.

### ACKNOWLEDGMENTS

We thank members of the Eyo lab, the Perez-Reyes lab, the University of Virginia Epilepsy Interest Group (EIG) and the Center for Brain Immunology and Glia (BIG) for valuable discussions in the development of this project. We acknowledge contributions from Dr. Kaushik Sharma in the initiation of this project, the gift of reagents from Dr. Tajie Harris and the generous provision of TRAP2 mice by Dr. Jaideep Kapur. Graphical illustrations were made using BioRender (https://biorender.com/). This work was supported by grants from The National Institutes of Health (R01NS122782; awarded to U.B.E. and E.P-R) a Translational Pilot Grant from the University of Virginia Brain Institute (awarded to U.B.E. and E.P-R), a Pharmacology T32 training grant (GM007055 awarded to S.G.) and funding from The Owens Family Foundation (awarded to U.B.E.).

## DATA AVAILABILITY STATEMENT

Data will be made available upon reasonable request to corresponding author as well as detailed protocol provided for any specific experimental procedures.

## FIGURES AND FIGURE LEGENDS

**SUPPLEMENTAL FIGURE 1.**
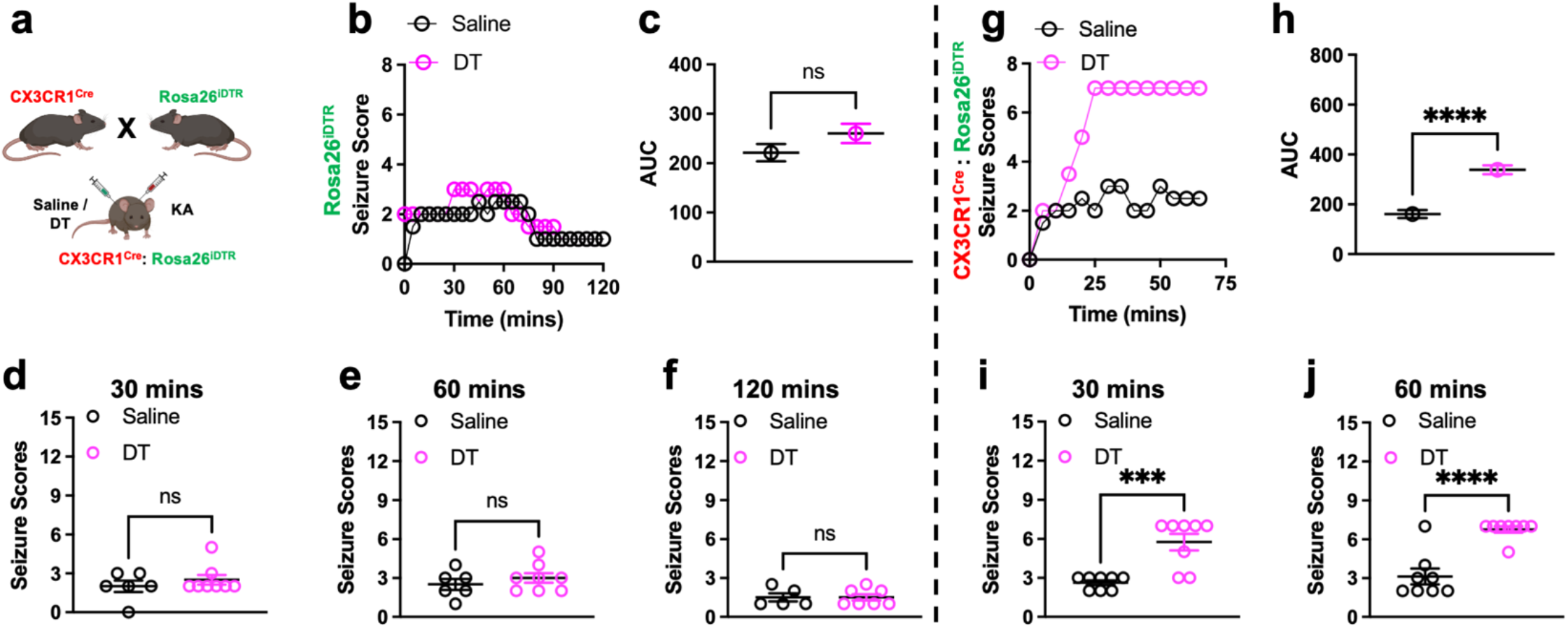
Pharmacogenetic microglial elimination and chemoconvulsive seizures. (a) Mating scheme of mice generation and experimental setup. (b and c) Overall (b) and average (c) Racine seizure scores from saline or diphtheria toxin (DT) treated Rosa26^iDTR^. n = 6-8 mice per group (d and e) Overall (d) and average (e) Racine seizure scores from saline or DT treated CX3CR1^Cre^: Rosa26^iDTR^ mice. n = 8 mice per group. Data presented as median in b and d and mean ± s.e.m in c and e. Statistics calculated by Student’s T-test, ****p < 0.0001.

**SUPPLEMENTAL FIGURE 2.**
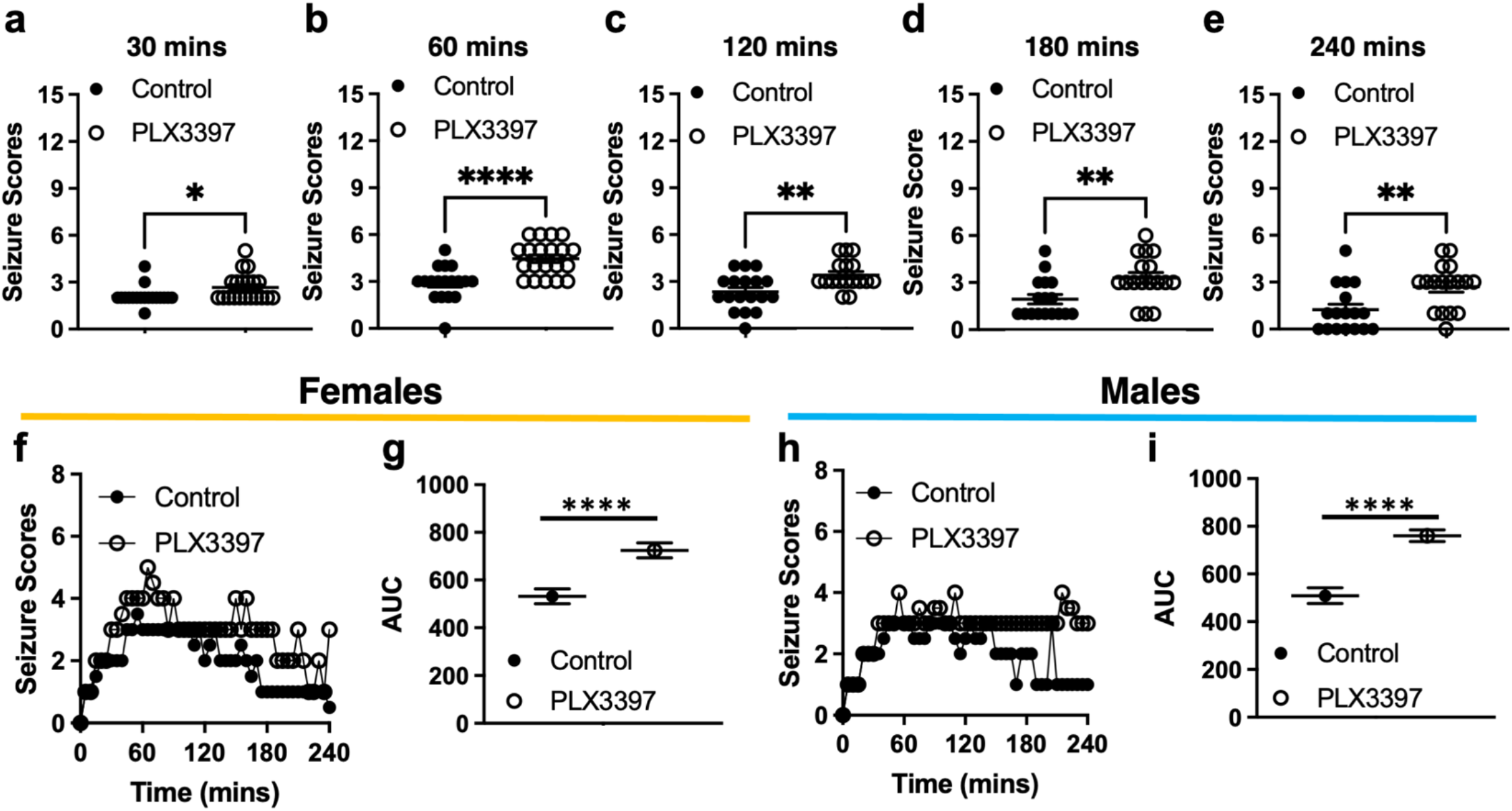
Time and sex features of chemoconvulsive kainic acid-induced seizures with microglial pharmacological elimination. (a- e) Racine seizure scores at various time points from control and PLX3397 treated mice over 240 mins. n = 18-20 mice per group. (f - i) Overall (f, h) and area under the curve (AUC, g, i) Racine seizure scores from control and PLX3397 treated female (f – g) and male (h – i) mic. Data presented as mean ± s.e.m in a – e, g and i and as median in f and h. Statistics calculated by Student’s T-test, *p < 0.05, **p < 0.01; ****p < 0.0001.

**SUPPLEMENTAL FIGURE 3.**
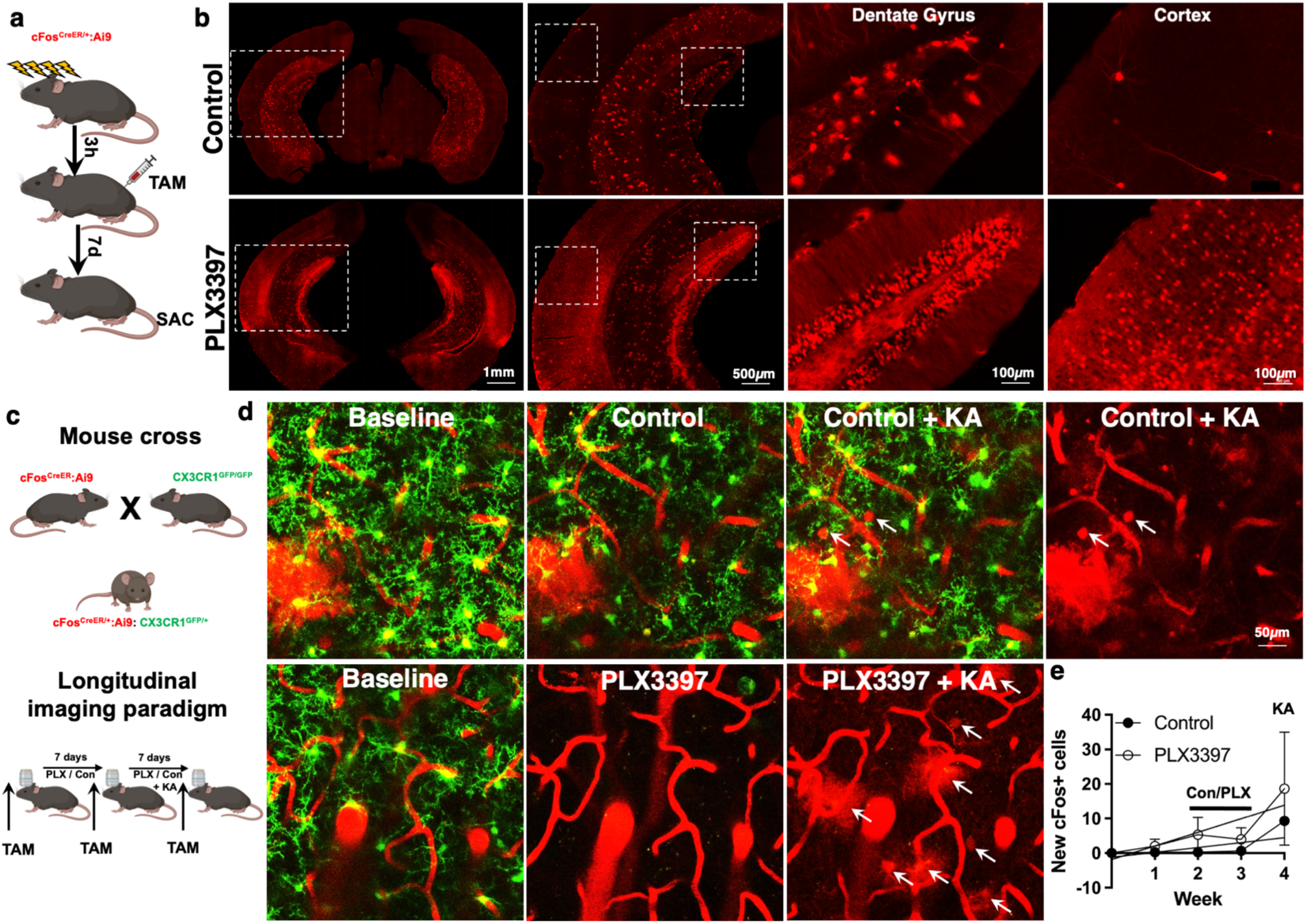
Pharmacological microglial elimination increases brain activity. (a) Experimental protocol for seizure cFos cell labelling following KA-induced seizures (b) Representative images of tdTomato-expression under the cFos promoter in cFos^tdTomato^ mice under baseline conditions and at 3h of KA after a 7-day treatment of control of PLX3397 in the cortex and hippocampus. (c) Experimental cross (top) and scheme for monitoring protocol for seizure cFos cell labelling following KA-induced seizures (bottom). (d) Representative longitudinal *in vivo* two-photon images of microglial (green), blood vessels (tubular red structures), and cFos positive (red circular structures) in different conditions. (e) Quantification of the average intensity of tdTomato fluorescence in the different conditions with longitudinal *in vivo* two-photon imaging. Mice were monitored for 2 weeks without PLX3397 and then for two weeks with control or PLX3397 chow before treatment with KA. n = 2 mice per group. Data presented as mean ± s.e.m.

**SUPPLEMENTAL FIGURE 4.**
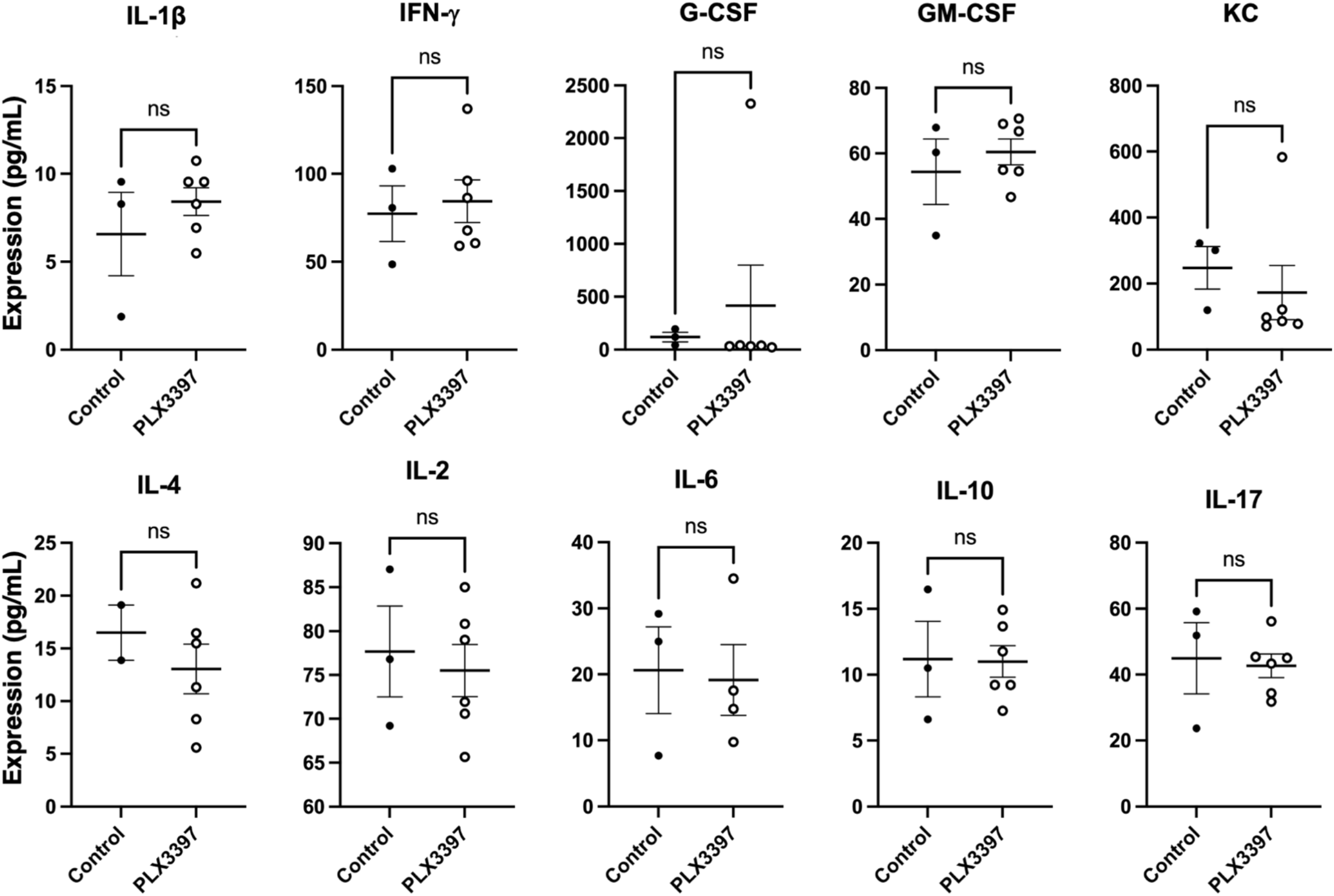
Microglial elimination does not alter cytokine levels. (Expression levels for various cytokines from control and PLX3397-treated brains after a week of treatment. n = 2-5 mice per group. Statistics calculated by Student’s T-test. Data presented as mean ± s.e.m.

**SUPPLEMENTAL FIGURE 5.**
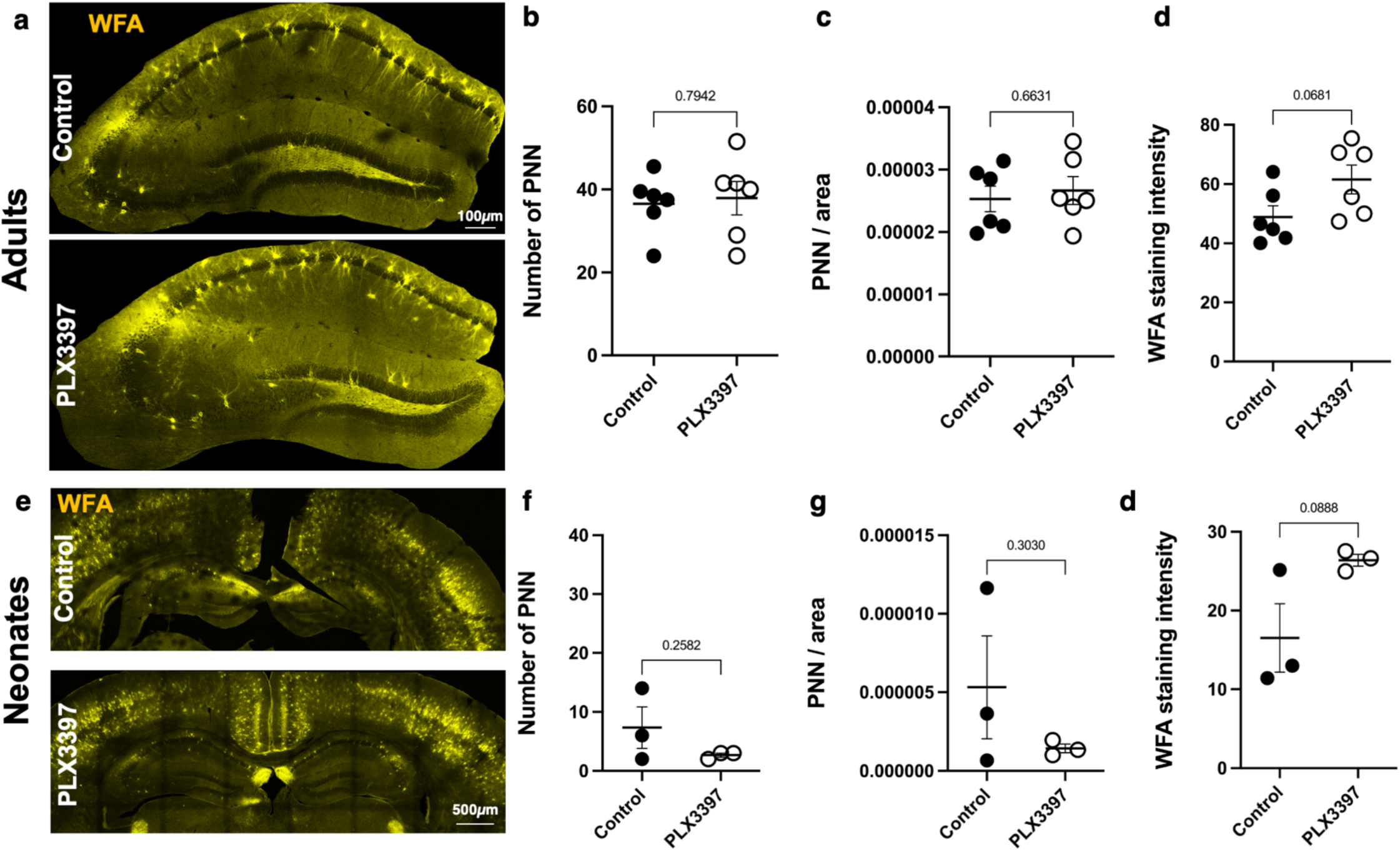
Microglial elimination does not alter perineuronal net density. (a) Representative images of hippocampal tissues stained with WFA (yellow) to label perineuronal nets (PNNs) from control and PLX3397-treated adult brains after a week of treatment. (b - d) Quantification of the number of PNN (WFA^+^) cells (b), the area occupied by PNN (WFA^+^) in hippocampal tissue (c), and the fluorescence intensity of WFA (d) from control and PLX3397-treated brains after a week of treatment. n = 6 mice per group. (e) Representative images of hippocampal tissues stained with WFA (yellow) to label perineuronal nets (PNNs) from control and PLX3397- treated neonatal brains after 2 days of treatment. (f - h) Quantification of the number of PNN (WFA^+^) cells (f), the area occupied by PNN (WFA^+^) in hippocampal tissue (g), and the fluorescence intensity of WFA (h) from control and PLX3397-treated brains after 2 days of treatment. n = 3 mice per group. Statistics calculated by Student’s T-test. Data presented as mean ± s.e.m.

**SUPPLEMENTAL VIDEO S1.**
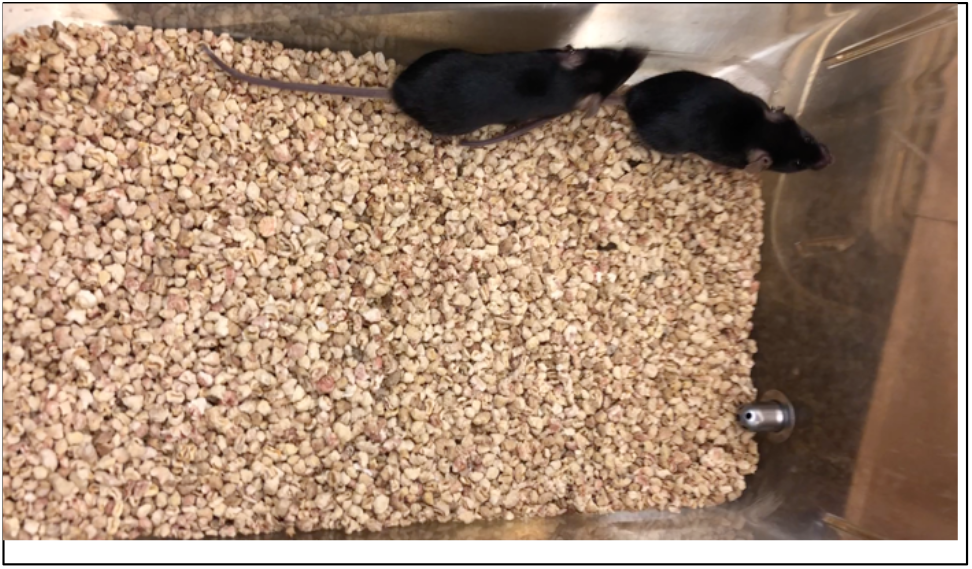
Mouse activity at 2 days of KA treatment in control mice. Wildtype mice exploring their cage at 2 days after experiencing KA-induced status epilepticus. Mice are actively exploring their environment. The movie is 15 seconds long.

**SUPPLEMENTAL VIDEO S2.**
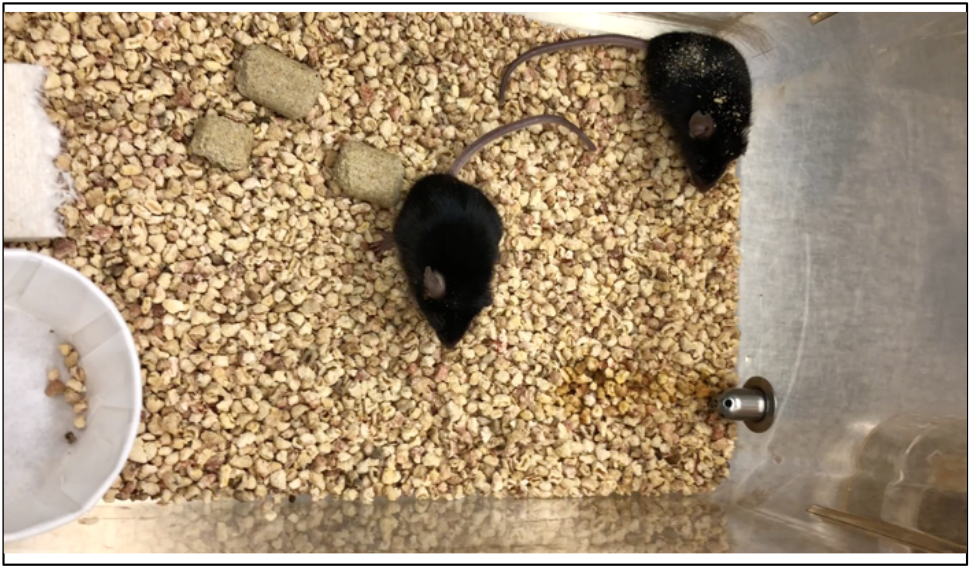
Mouse activity at 2 days of KA treatment in PLX3397-treated mice. Wildtype mice were exposed to PLX3397 for 7 days and then treated with KA. This movie is collected at 2 days after KA-induced status epilepticus and mice are not as active in exploring their cage. The movie is 16 seconds long.

**SUPPLEMENTAL VIDEO S3.**
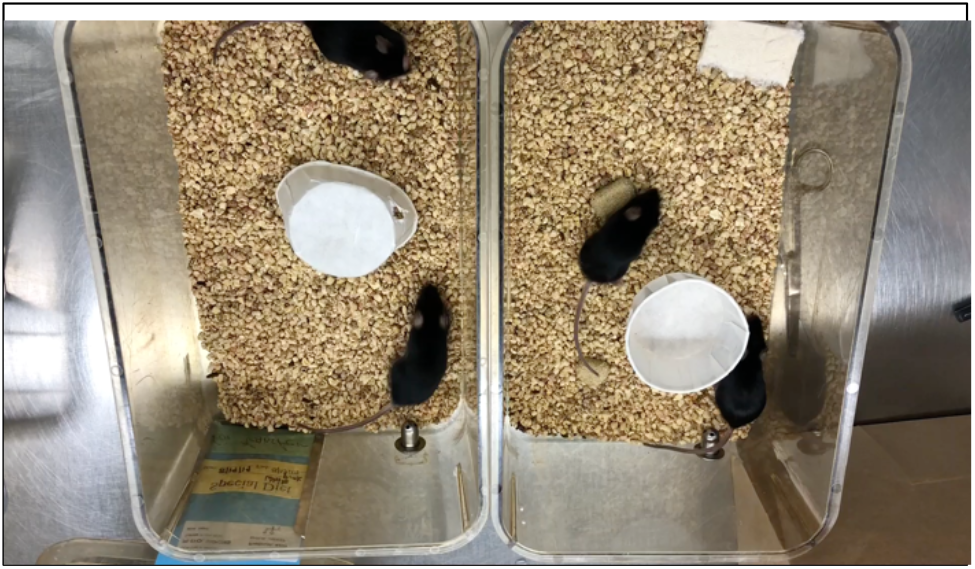
Mouse activity at 3 days of KA treatment in control and PLX3397-treated mice.Wildtype mice were exposed to either control (left) or PLX3397 (right) chow for 7 days and then treated with KA. This movie is collected at 3 days after KA-induced status epilepticus and PLX3397-treated mice are not as active in exploring their cage The movie is 31 seconds long.

**SUPPLEMENTAL VIDEO S4.**
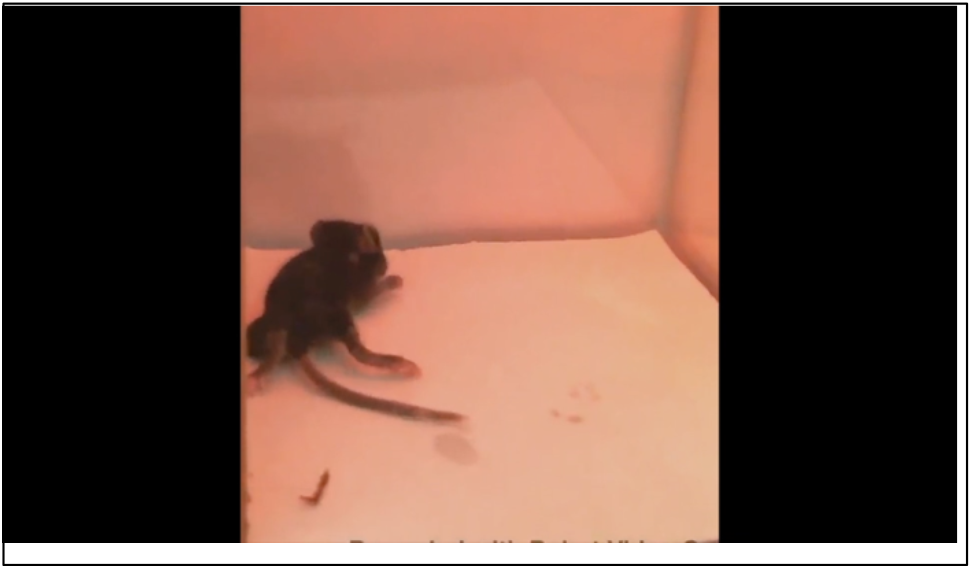
Neonatal mouse undergoing febrile status epilepticus. Neonatal p14 mouse exposed to hyperthermia showing whole body convulsions. The movie is 1 minute long.

